# Decoding network-mediated retinal response to electrical stimulation: implications for fidelity of prosthetic vision

**DOI:** 10.1101/2020.06.29.178723

**Authors:** Elton Ho, Alex Shmakov, Daniel Palanker

## Abstract

**Objective:** Patients with the photovoltaic subretinal implant PRIMA demonstrated letter acuity by ~0.1 logMAR worse than the sampling limit for 100μm pixels (1.3 logMAR) and performed slower than healthy subjects, which exceeded the sampling limit at equivalently pixelated images by ~0.2 logMAR. To explore the underlying differences between the natural and prosthetic vision, we compare the fidelity of the retinal response to visual and subretinal electrical stimulation through single-cell modeling and ensemble decoding.

**Approach:** Responses of the retinal ganglion cells (RGC) to optical or electrical (1mm diameter arrays, 75μm pixels) white noise stimulation in healthy and degenerate rat retinas were recorded via MEA. Each RGC was fit with linear-non-linear (LN) and convolutional neural network (CNN) models. To characterize RGC noise level, we compared statistics of the spike-triggered average (STA) in RGCs responding to electrical or visual stimulation of healthy and degenerate retinas. At the population level, we constructed a linear decoder to determine the certainty with which the ensemble of RGCs can support the *N*-way discrimination tasks.

**Main results:** Although LN and CNN models can match the natural visual responses pretty well (correlation ~0.6), they fit significantly worse to spike timings elicited by electrical stimulation of the healthy retina (correlation ~0.15). In the degenerate retina, response to electrical stimulation is equally bad. The signal-to-noise ratio of electrical STAs in degenerate retinas matched that of the natural responses when 78±6.5% of the spikes were replaced with random timing. However, the noise in RGC responses contributed minimally to errors in the ensemble decoding. The determining factor in accuracy of decoding was the number of responding cells. To compensate for fewer responding cells under electrical stimulation than in natural vision, larger number of presentations of the same stimulus are required to deliver sufficient information for image decoding.

**Significance:** Slower than natural pattern identification by patients with the PRIMA implant may be explained by the lower number of electrically activated cells than in natural vision, which is compensated by a larger number of the stimulus presentations.

## Introduction

Age-related macular degeneration (AMD) is a leading cause of untreatable blindness. Geographic atrophy (GA), the atrophic form of advanced AMD, affects around 3% of people above the age of 75, and around 25% - above 90 (1, 2). Due to gradual loss of photoreceptors in the central macula, GA patients experience severe deterioration in high-resolution central vision, compromising their ability to read and recognize faces. Although central vision degrades over time, patients retain their low-resolution peripheral vision, and hence typically do not lose visual acuity beyond 20/400.

In healthy retina, optical information (local light intensity) is converted via phototransduction into decrease of the cell potential (hyperpolarization) in photoreceptors. Rods are responsible for monochromatic vision at low light intensities, while cones, operating in brighter light, provide color vision at high resolution. Decrease in cell potential reduces the rate of release of neurotransmitter glutamate in synapses with the secondary neurons – bipolar and horizontal cells. By providing lateral inhibition that forms an antagonistic surround, horizontal cells perform the first step in spatial contrast enhancement. OFF bipolar cells receive their input from photoreceptors via sign-preserving ionotropic synapses and hence respond to light stimuli by hyperpolarization. ON bipolar cells receive their input via sign-inverting metabotropic synapses, resulting in depolarization in response to light. Bipolar cells electrically integrate inputs from multiple photoreceptors and relay these signals to tertiary retinal neurons – amacrine and ganglion cells (RGCs). Amacrine cells regulate the inputs into the RGCs mostly through inhibition. Finally, RGCs digitize these signals into bursts of the action potentials (“spikes”), which propagate via optic nerve to the brain. Different types of RGCs encode visual information via various mechanisms and properties: ON and OFF pathways, sizes of receptive fields, transient and sustained responses, chromatic sensitivity, etc. Signals from the overlapping mosaics of the various types of RGCs are further processed in the brain before merging into a single visual percept.

In atrophic AMD, photoreceptors in the central macula slowly degenerate and disappear, while the inner retinal neurons remain largely intact, albeit with some rewiring (3–5). Since horizontal cells connect to the terminals of photoreceptors, in geographic atrophy they become detached from the remaining neural network.

One approach to restoration of sight in retinal degeneration is to replace the missing photoreceptors with photodiodes (6, 7), which convert the incident light into electric current flowing through the retina, and thus convey visual information to the secondary neurons by electrical stimulation. Massive amplification mechanisms in photoreceptors enable their operation in a very broad (about 10 orders of magnitude) range of light intensity. Photodiodes require much brighter illumination (about 1 mW/mm^2^) in order to provide sufficient current for retinal stimulation. Therefore, photovoltaic system for restoration of sight includes augmented-reality glasses, where images captured by the camera are projected into the eye using more intense light. To avoid perception of this intense light by the remaining photoreceptors, near-infrared wavelength (880nm) is used. To provide charge-balanced electrical stimulation, the light is pulsed, and to enable stable visual percepts, pulse repetition rate should exceed the frequency of flicker fusion (around 30 Hz).

Ex-vivo and in-vivo animal studies with this system demonstrated preservation of multiple features of the natural retinal signal processing: sizes of the RGC receptive fields, with antagonistic surround (8) and non-linear summation of subunits (6), flicker fusion (9) and adaptation to static images (10), and spatial resolution matching the pixel pitch (55 and 75 μm) (9). Interestingly, both, ex-vivo and in-vivo studies demonstrated OFF responses. This may be explained by stimulation of rod bipolar cells, which feed into the ON and OFF cone pathways via amacrine cells. Contrast sensitivity of prosthetic vision in rats appears to be about 5 times lower than natural (11), which could be partially compensated by image processing prior to projection into the eye. Clinical trial demonstrated that patients correctly perceive various patterns of lines and letters, demonstrating monochromatic shaped vision with resolution closely matching the pixel size (100 μm in the first trial) (7). They also report flicker fusion at frequencies exceeding 30 Hz.

One of the features of such prosthetic vision, however, appears to be the lower than normal speed of the pattern and letter recognition. In the first trial, it took about 4 seconds for letter identification by patients with PRIMA implants (7), while it takes less than half a second in normal subjects, when font sizes exceed the acuity limit (12). Here we investigate the potential retinal underpinnings of this phenomenon by comparing RGC responses to visual and electrical stimulation in healthy and degenerate rat retina recorded on a multi-electrode array (MEA). In particular, we assess the amount of noise in various cellular responses, as well as the strategies for image recognition based on ensemble encoding by a population of cells.

## Methods

### Photovoltaic implants

Photovoltaic arrays (1mm diameter, 30 μm in thickness, with 75 μm pixels) (Fig. 1a) were manufactured from crystalline silicon, as described earlier (13), to produce anodic-first pulses. Active electrode was 20 μm in diameter, and each pixel was surrounded by a return electrode, connected into a mesh common to all pixels. Both electrodes were coated with SIROF film of about 300 nm in thickness (9).

**Figure 1.**
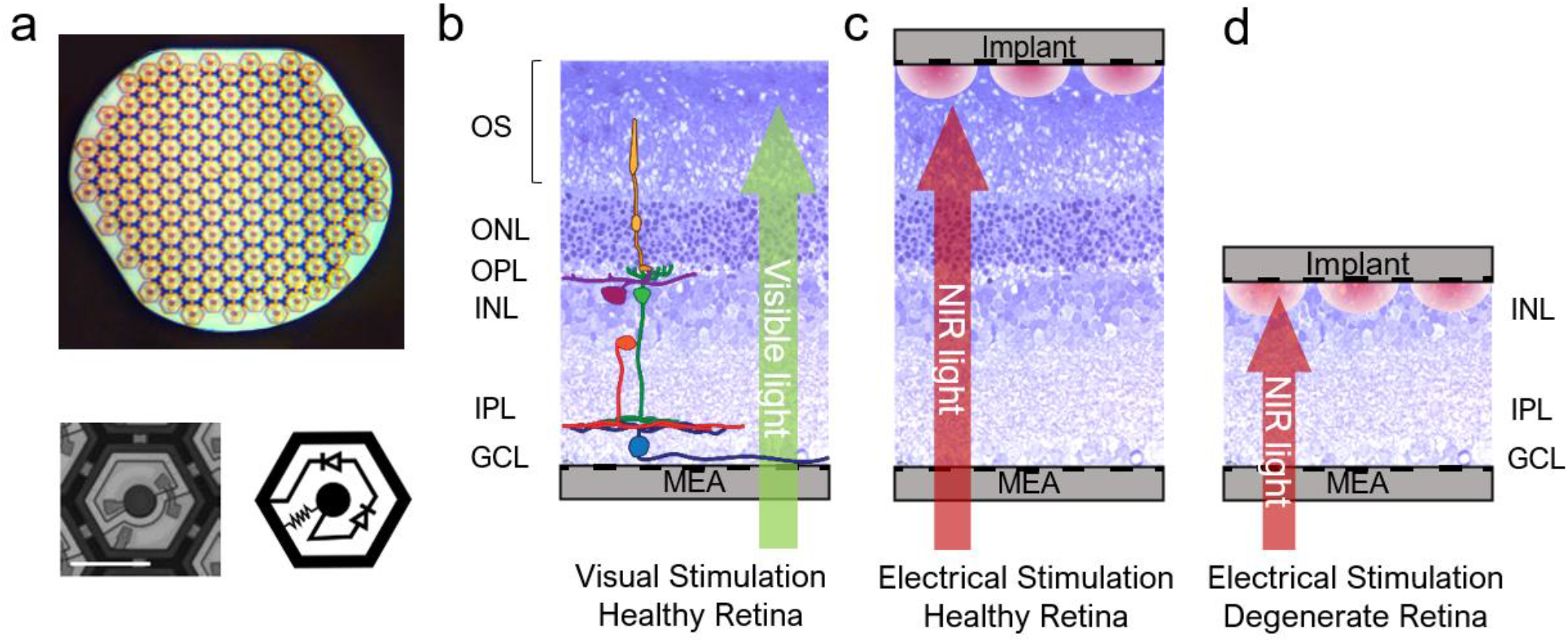
Photovoltaic subretinal implant and stimulation scheme. (a) Top: implant of 1mm in width with 70μm pixels. Bottom left: a single photovoltaic pixel. Bottom right: circuit diagram of a pixel. (b) Visual stimulation of the healthy rat retina placed on top of a transparent MEA. (c) Electrical stimulation of the healthy retina by an implant placed on top of photoreceptors. (d) Electrical stimulation of the degenerate retina by an implant placed on top of the inner nuclear layer (INL).

### Retinal recording

We used Long-Evans (LE, n=4) and Royal College of Surgeons (RCS, n=4) rats for healthy and degenerate retinal models, respectively. Eyes were enucleated from euthanized (390mg/kg pentobarbital sodium, 50mg/mL phenytoin sodium) rats. A section of the retina (~3mm × 3mm) was dissected and placed ganglion cells side facing a 512-electrode multielectrode array (MEA) (Fig. 1b) (14). The retina was constantly perfused with Ames’ medium at 29.4 °C and bubbled with a mixture of 95% O_2_ and 5% CO_2_. For electrical stimulation, an implant was placed onto the subretinal side of the tissue (Fig. 1c, d). A nylon mesh (~100μm cell size) was used to press the implant and retina onto the MEA for better contact (10). Voltage waveforms from each of the 512 electrodes on the MEA were sampled at 20 kHz frequency, amplified and digitized using custom-made readout electronics and data acquisition system (14).

### Stimulation protocol

For electrical stimulation, an 880nm diode laser coupled via a 400μm multimode fiber was used for illumination. The beam exiting from the fiber was collimated and homogenized using a 2^°^ divergence microlens array. In the same optical path, we placed a yellow LED (591nm) for visual stimulation. Both light sources were used as backlighting for an LCD screen (Holoeye HEO-0017) to generate images (6, 10). The 8-bit LCD panel had a 60Hz native frame rate, 1024×768 resolution, and a white-to-black intensity ratio of 10000:1 at 591nm and 200:1 at 880 nm. Projected onto the retina, each screen pixel formed a 6 × 6 μm^2^ square.

To characterize spatiotemporal properties of RGCs, a spatiotemporal binary white noise stimulus was used, where each pixel in each frame had a 50% chance of being bright or dark (15). The white noise for visual stimulation was shown at 30Hz frame rate, and made up of pixels of 60μm in width on the retina. The white noise for electrical stimulation was displayed at 20Hz frame rate, with the backlight laser pulsing at 4ms, and consisted of pixels of 70 μm in width on the retina. Each white noise stimulus lasted for 30 minutes.

### Spike sorting

Electrical stimulation generated artifacts by saturating the recording amplifiers, so part of the recorded waveforms had to either be discarded or adjusted. The recording of the first 8.25ms after the laser pulse was replaced with a randomly generated noise (“blanking”) that matched the noise level of the electrode. All action potentials during this period were lost, which may lead to underestimation of the cell responsiveness. Afterwards, to remove any lingering capacitive decay outside of the blanked period, we fitted the trace with a 7^th^-order polynomial, and then subtracted it out from the original trace.

The artifact-subtracted raw data were then used to find and sort the action potentials (“spikes”). A negative voltage deflection exceeding three times the root-mean-squared noise on each electrode was considered a spike. Custom-made software was used to perform spike sorting, as described previously (6, 14, 16). We applied dimensionality reduction to the detected spike waveforms using a principle component analysis, followed by expectation-maximization clustering (14). For each putative neuron, we calculated its electrophysiological image (EI), which is the average electrical signal measured on the whole MEA when the neuron produced a spike. An EI typically shows the soma location and axonal trajectory of the RGC (17) (18). For our analysis, we only included cells that responded to electrical stimulation, and have their somas completely or partially under the implant. Figures 2a and b illustrate examples of the populations of included cells for healthy and degenerate retinas, respectively. Figures 2c-e show example receptive fields and time courses (spike-triggered average, STA) for natural visual response, electrical stimulation of the healthy retina, and electrical stimulation of RCS retina, respectively. Electrically stimulated cells generally have faster, but weaker STAs, similar to previous observations (8, 11).

**Figure 2.**
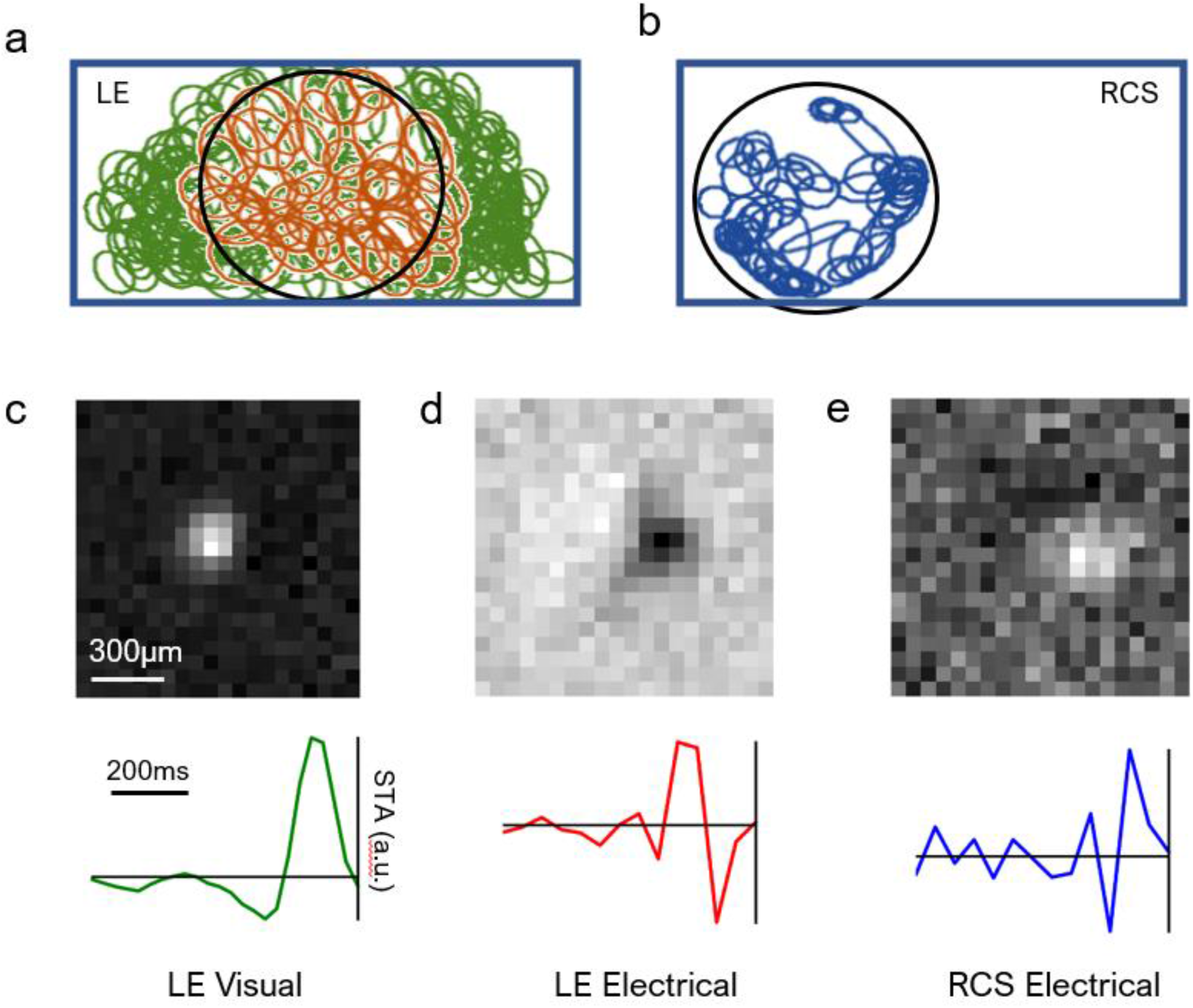
MEA recording of the RGC responses to stimulation. (a) Mosaic of the natural receptive fields (RF) in LE retina. The 1mm × 2mm rectangle marks the boundary of the MEA, and the 1 mm circle indicates the edge of the implant. Red ellipses correspond to cells responding to both visual and electrical stimulation, and green ellipses correspond to cells responding only to visual stimulation. Only the red cells were included in our subsequent analyses. (b) Mosaic of receptive fields in RCS retina upon electrical stimulation. (c) RF and the time course (STA) of a cell in the LE retina responding to visual white noise. (d) Same cell responding to electrical stimulation. (e) RF and the time course of a cell in the RCS retina responding to electrical stimulation.

### Modeling the RGC responses

Experimental data for each cell was fitted to a linear-nonlinear (LN) model (15) and a convolutional neural network (CNN) model (19), as illustrated in Fig. 3. Due to drifting in recording data, we used a train-test-discard split of 20/20/60 (see Supp. Materials). When comparing the model predictions to experimental data, we applied Gaussian broadening to each spike with *σ* = 2 white noise frames for smoothening (Fig. 5a). We then computed the Pearson correlation coefficient for the resulting traces. In addition, all model fits were 5-fold cross-validated, where we equally sized the sections dividing the entire recorded data. Neither model over-fitted to any particular segment of the data.

**Figure 3.**
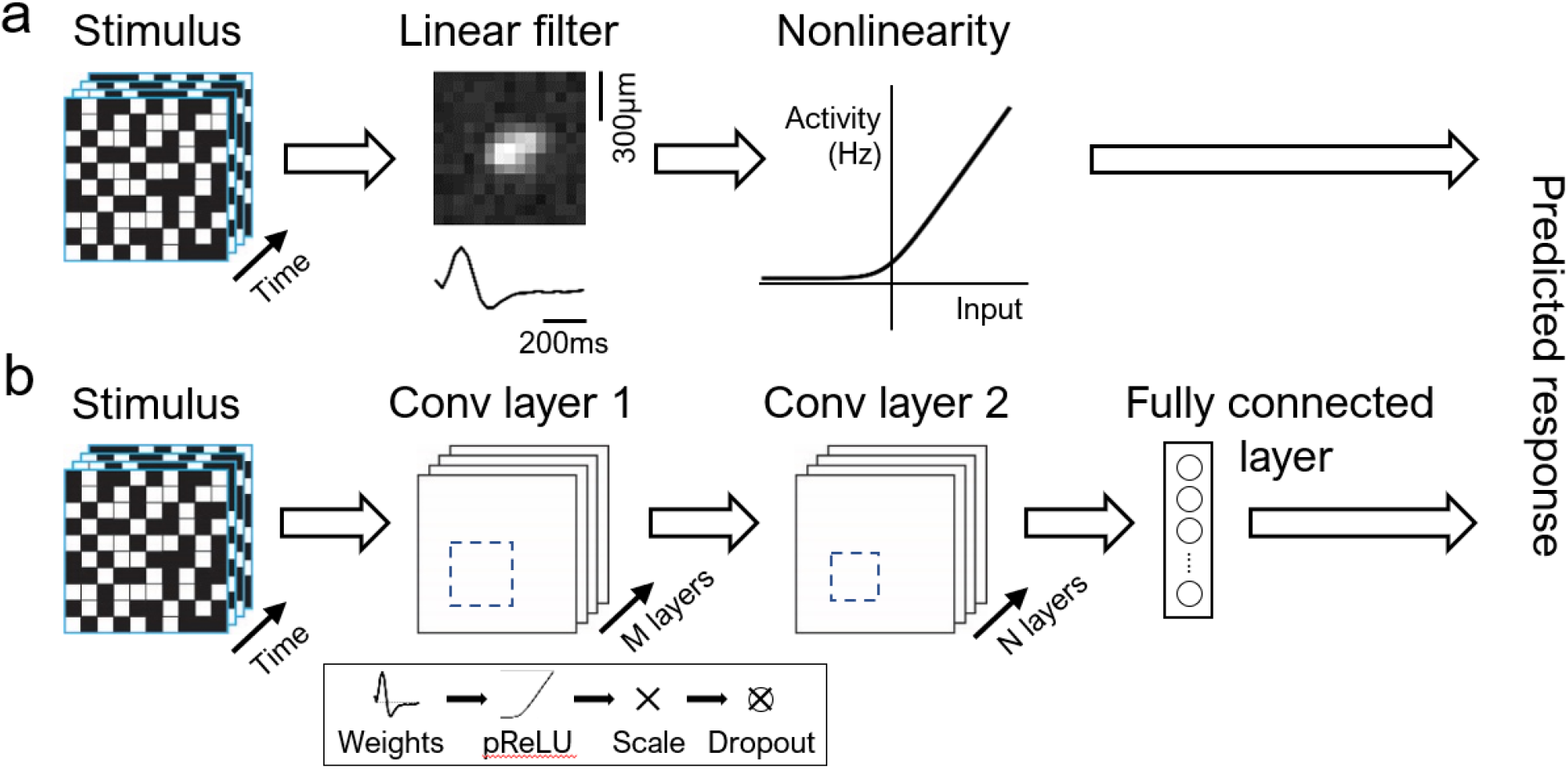
Computational models for individual RGCs. (a) Linear-nonlinear model. (b) Convolutional neural network. Each convolutional layer consists of a sequence of linear filters (weights), parametric rectifying linear unit (pReLU), batch renormalization (scaling), and a dropout layer.

#### LN model

Mathematically, the LN model (Fig. 3a) is set as follows:

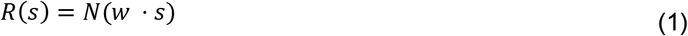

where *R* = response

*s* = stimulus

*w* = linear weights/filter

*N* = static nonlinearity

The linear filter *w* can be computed through STA response to the white noise stimulus. The static nonlinearity can be extracted by mapping the empirical cell activity to stimulus convoluted with the linear weights (*w* · *s*) (15).

#### CNN model

An implementation of a CNN model has been proven useful for modeling the healthy salamander retina (19), and here we used a similar architecture (Fig. 3b) with two convolution blocks followed by a dense layer. Each convolution block consisted of a two-dimensional convolution (weights), a parametric rectifier linear unit (pReLU), batch normalization (norm), and a dropout layer. These last two components sped up the training while also regularizing the network to prevent over-fitting. The number of filters and their dimensions for each convolution block are listed in Table 1. The output of the second conv layer was flattened into a one-dimensional vector before being fed into the final dense linear layer, which had a number of units matching the number of recruited RGCs in the retina.

**Table 1.** Number of filters and their dimensions for each convolution block and for each stimulation type.

|  |  | Stimulation |  |
| --- | --- | --- | --- |
|  |  | Visual | Electrical |
| Layer | Convolution block 1 | 8 filters, 13 x 13 each | 16 filters, 5 x 5 each |
|  | Convolution block 2 | 16 filters, 9 x 9 each | 32 filters, 5 x 5 each |

The network was trained using the gradient-descent ADAM optimizer (20) and a Poisson log-likelihood. L2 weight regularization was employed on the convolution and linear layers, while L1 regularization was used on the output of the network. Especially for cells with lower firing rates, L1 can efficiently zero-out many weights. The complete loss function was defined as follows:

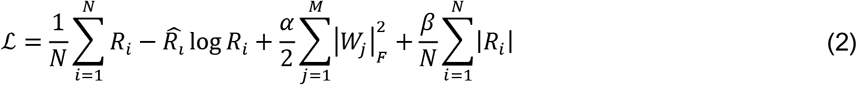

where *R*_*i*_ = model response *i*

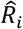 = target response *i*

|*W*_*j*_|_*F*_ = Frobenius norm of the *j-th* convolution or linear layer weight matrix

*N* = total number of training samples

*M* = total number of weight matrices

α, β = L2 and L1 regularization coefficients, respectively.

The input stimulus was similar to that used in computing STAs, while the response now included all activity and inactivity. For visual stimulation, 20 consecutive movie frames (spanning 600ms) were considered one stimulus, and the spike rate during 33ms following the stimulus was taken as the target response. Similarly, for electrical stimulation, 5 movie frames (250ms) and the following 50ms of activity was considered a stimulus-response pair. To improve precision of spike timing while increasing the training sample count, the electrical stimulus was up-sampled with linear interpolation to 250Hz, and the corresponding RGC spiking activity was binned to match the stimulus frame rate. During validation, the predicted activations were down-sampled back to the original frame rate before the correlation was computed.

The CNN is parameterized by 13 different hyperparameters, including filter count, size, stride, and nonlinearity for each of the two convolution blocks. In addition, we also explored different values for the learning rate, L1 and L2 coefficients, batch size, and dropout probability. We performed 100 trials for each dataset with randomized values for all parameters using the SHERPA hyperparameter optimization library (21). Networks were trained for 50 epochs on electrical datasets and 50 epochs on visual datasets. The total training time on a single Nvidia Titan × GPU was 30 minutes and 2 hours, respectively.

### RGC noise estimation

We estimated the noise level in RGC firing under electrical stimulation using the algorithm illustrated in Fig. 4a. First, we passed the white noise stimulus through a LN model for an LE RGC under visual stimulation, generating a simulated spike train. We then removed spikes randomly at some predefined noise ratio (NR). Spontaneous spikes were added into the spike train to match the original average spike rate via a Poisson process. With the new noise-injected spike train, we re-computed the STA. The boxed regions in Fig. 4b with dimensions (length, width, time) = (3 px, 3 px, 4 frames) were used for the subsequent analyses.

**Figure 4.**
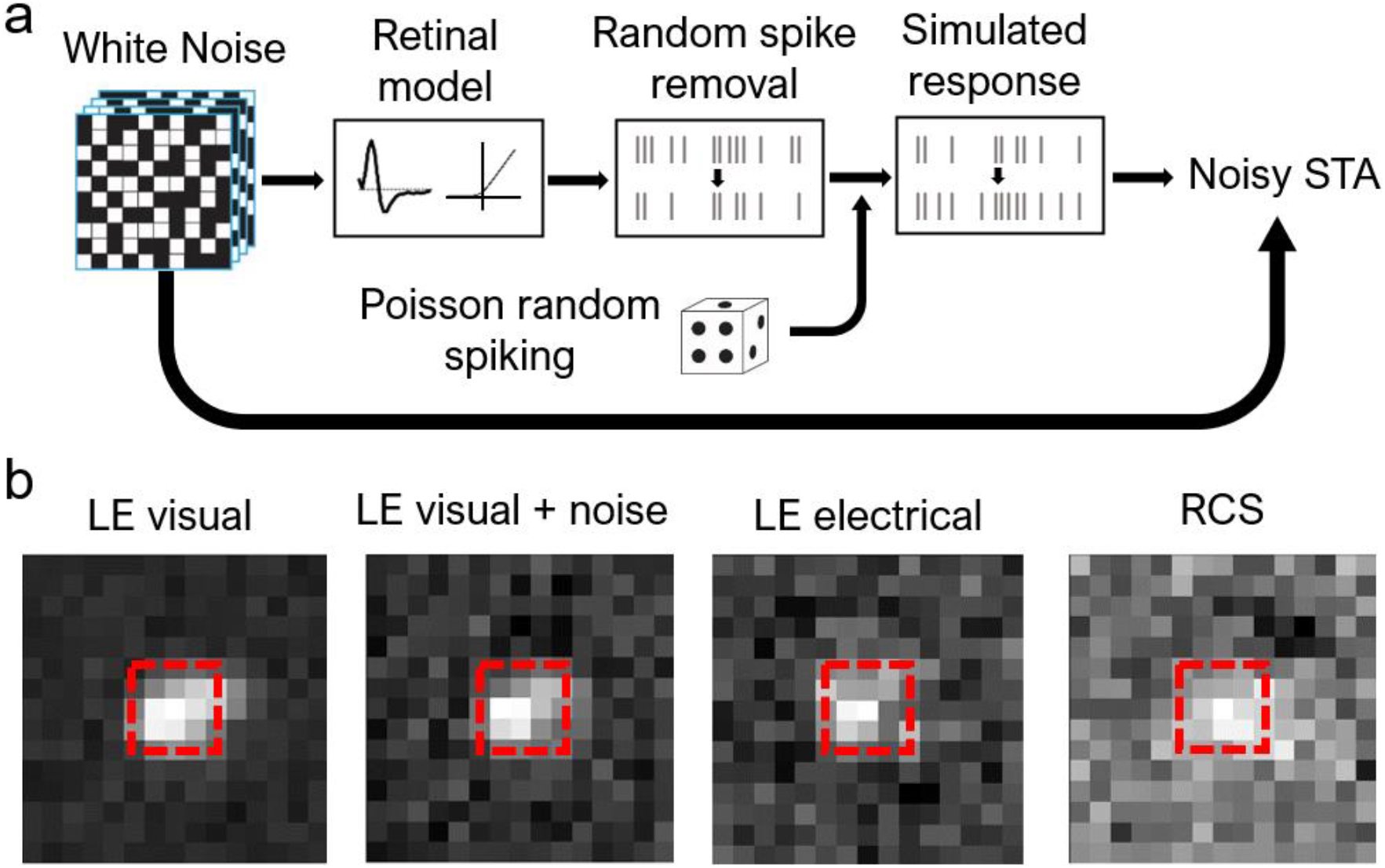
Adding noise to STAs. (a) Algorithm used for adding noise to STAs. Spikes were first generated by a model of the visual response, and then partially removed and replaced with spikes generated through a Poisson process. The resulting simulated response was then used to re-compute the STA. (b) Examples of STAs under different stimulation and retina types. With properly chosen noise ratio, the noise-injected visual STA resembles that of the LE retina under electrical stimulation. The region bound by the red dash line was used for subsequent analyses.

**Figure 5.**
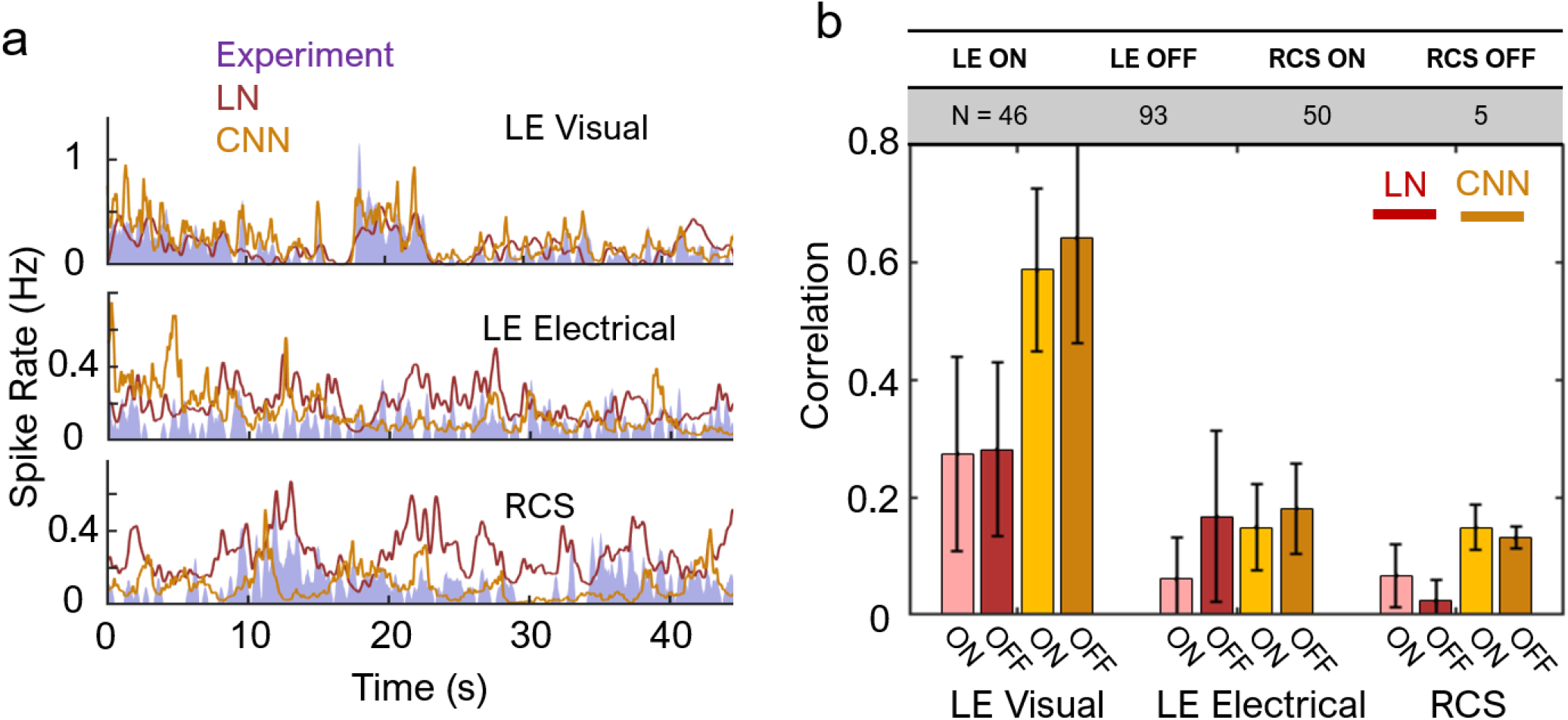
Single-cell model predictions and its accuracy. (a) Examples of LN and CNN model predictions for a single cell, alongside the experimental data. Both models fit well the LE visual responses, but not the electrical stimulation of either LE or RCS retinas. (b) Correlation with the test set data, averaged over a population of cells (table). Model fit to electrically stimulated cells (~0.15 for CNN with LE and RCS data) is significantly worse than to the visual response (~0.6).

For each cell and its STA, we computed a characteristic correlation curve (CCC) and its corresponding area-under-curve (AUC), as shown in Fig. 6a. Let *F*(*n*) be the 24 STA frames preceding the n-th spike, *N* be the total number of spikes, and 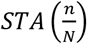 be the STA computed with only a fraction *n*/*N* of the spike train. We have

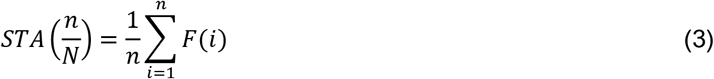

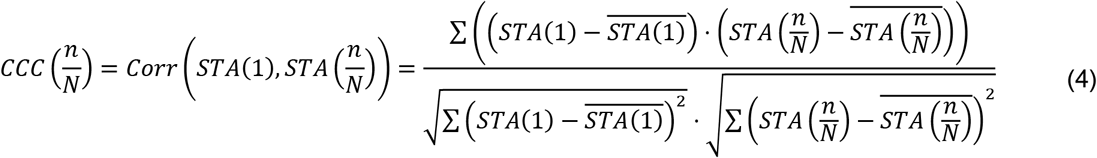

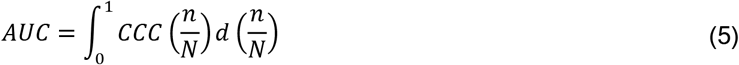

**Figure 6.**
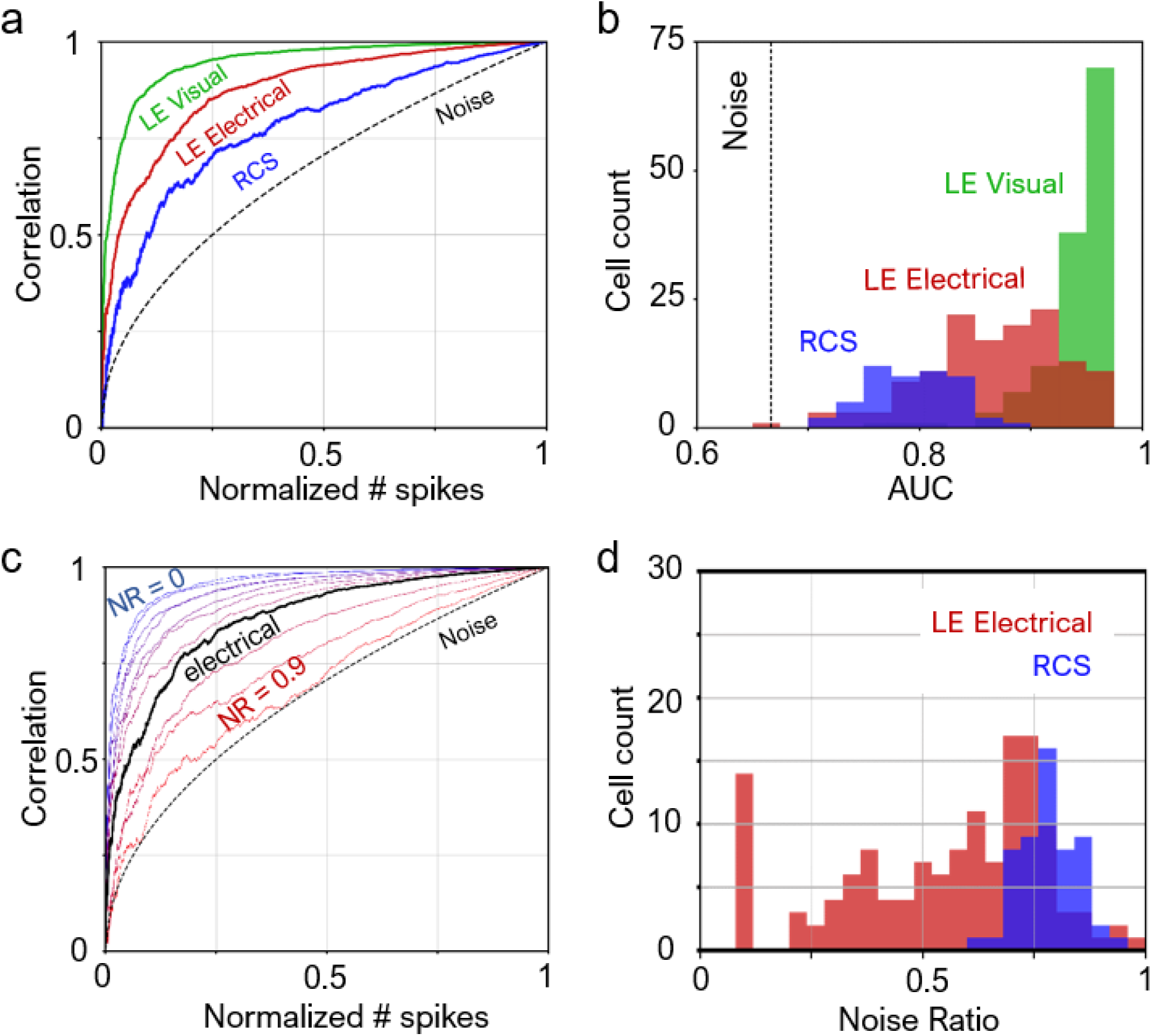
Noise in retinal responses. (a) Characteristic correlation curves. A cell with little noise would have a curve farther away from the pure noise curve. As a corollary, the greater the area-under-curve (AUC), the less noisy is the cell. (b) AUC for a population of cells. Almost all LE cells responding to visual stimulus exhibit less noise than all RCS cells under electrical stimulation. (c) Example correlation curves for LE visual with different added noise ratios (NR). At NR ~ 0.65, the noise-injected correlation curve matches that for LE electrical. (d) Noise ratios over the cell population. For LE electrical, the average noise ratio is 55.3±22.5%; for RCS, the noise ratio is 78.2±6.5%.

Note that *STA*(1) is when all the spikes are included in computing the STA.

If a cell responds perfectly only to one single type of stimulus with no spontaneous firing, then 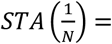 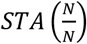, and AUC = 1. Figure 6a shows the CCC’s of three example cells, and Fig. 6b shows the distribution of cell count for various levels of AUC.

Out of all LE retinas, we selected the cell that had the median AUC under visual stimulation as a reference. Noise-injection into the STA of this cell yielded a family of CCCs shown in Fig. 6c. To characterize the noise of each cell under electrical stimulation, we matched its CCC to the curve in the family that has the most similar AUC, and the resulting matching noise ratio characterizes the cell (Fig. 6d).

### Ensemble encoding

To evaluate how much information is encoded by the ensemble of cells for the pattern recognition task, we simulated projection of pixelated Landolt-C onto a piece of retina (Fig. 7a). Each presentation of the C lasted for 5 movie frames, and was spatially pixelized into either 70 μm (for electrical stimulus) or 60 μm (for visual stimulus) pixels. The brightness of each pixel was then rounded to the nearest one of 8 evenly spaced greyscale levels. The resulting simulated stimulus 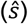 had dimensions (length, width, time) = (64, 32, 24 frames) for visual stimuli or (20, 20, 24 frames) for electrical stimuli. The first 19 frames were all dark, and the remaining frames were bright, where the Landolt-C was displayed. The simulated stimulus was then convoluted with the STAs (*w*) of each cell to produce an input strength 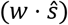, similar to Eq. (1). We then inspected the white noise stimulus for blocks of consecutive frames (*s*, same dimensions to 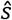) that share a similar input strength, mathematically defined as

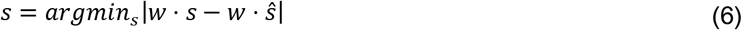

**Figure 7.**
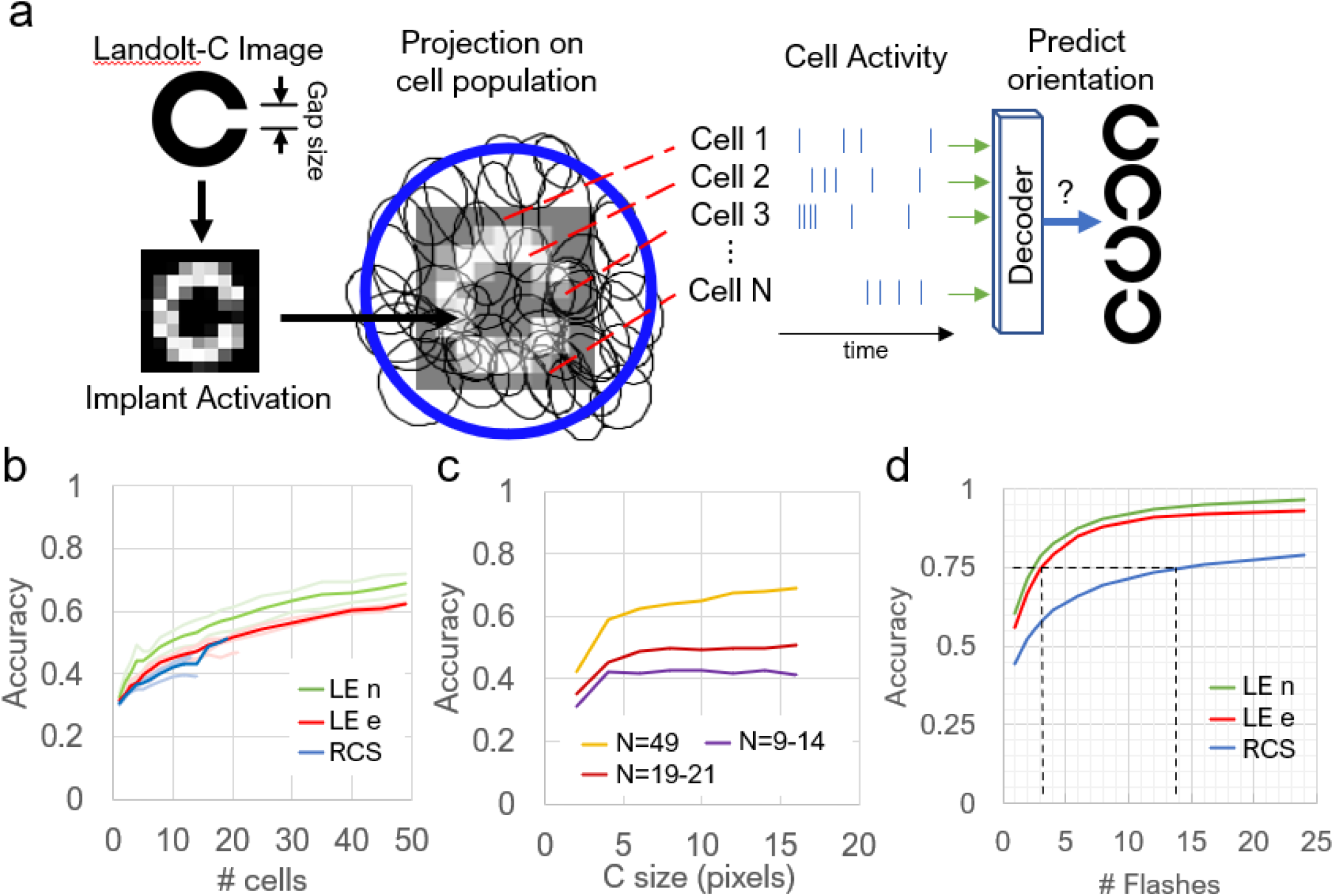
Ensemble encoding of the visual information. (a) Algorithm for evaluation of the ensemble encoding accuracy. Using the cell activity under a certain projection of a Landolt-C, decoder discerns its orientation. (b) Accuracy versus number of cells included in the decoder, with a C size of 14 pixels. Faint lines represent individual trials. (c) Accuracy as a function of the letter C size. (d) Accuracy versus number of presentations (flashes) of the letter. Since electrical stimulation activates fewer cells in the RCS retina than visual stimulation in healthy retina, ~5 times more flashes are required to achieve the same decoding accuracy as natural. (4 LE retinas, N cells = 20, 21, 49, 49; 4 RCS retinas, N cells = 9, 10, 13, 14)

The 500 blocks that best satisfy the above criterion are chosen, and the average RGC activity 30ms following each block was considered a response to the Landolt-C. Afterwards, responses of all cells were concatenated into a template with time bins of 5ms. Four different templates were created for four orientations (up, down, left, right) of the C using the same procedure. We then simulated 10,000 trials with random orientations. For each trial, the number of spikes in each time bin was simulated as a Poisson process with its mean matching the spike rate in the bin in the corresponding template. The generated spiking pattern for each trial was correlated to all templates, and one with the highest Pearson’s *r* was considered the decoded orientation. Decoding accuracy was taken as the ratio of correctly decoded trials to total. For the 4 LE retinas, the cell counts were 49, 49, 21, and 20; for the 4 RCS retinas, the cell counts were 19, 14, 13, and 9. To study the effect of number of cells on decoding accuracy, we fixed the size of the C at 14 pixels. To study the effect of C size and number of flashes, we included all cells on each retina into the decoder.

## Results

### Single-Cell response modeling

For the natural response of healthy retina, the LN model fitted to levels similar to previous reports in salamander and rat retinas (correlation in the range of 0.3) (19, 22). The CNN model fitted much better to the spike trains elicited by visual stimulation in ON and OFF cells (correlation of about 0.6, Fig. 5a top), agreeing with earlier studies in the salamander retina (23). However, both models predicted retinal responses to electrical stimulation of the healthy or degenerate retina significantly worse (Fig. 5a, center and bottom). Across a population of cells and multiple retinas (n=4 each), CNN fits to electrical data reached a correlation of only ~0.15, significantly lower than 0.6 for the LE visual response (Fig. 5b). The LN model fitted distinctly better to electrical OFF cells than ON cells in LE retinas (*p*<10^−7^, two-sample *t*-test), while the CNN model fitted with less discrepancy between the two cell types (*p*=0.013). For electrical ON cells, the CNN model fitted significantly better than the LN model (*p*<10^−9^), but the same cannot be said for electrical OFF cells. In RCS retinas, correlation with the CNN model was similar to the LE retina under electrical stimulation (Fig. 5b). However, correlations with the LN model were far worse for RCS OFF cells than for LE electrical OFF cell, while that for electrical ON cells remained similar.

### Noise estimation in RGC firing

Figure 6a illustrates the cross-correlation curves (CCC) of three example cells, as well as the CCC for an STA generated from randomly sampled white noise frames, which is described by a square root dependence on the normalized number of spikes. The CCC for RCS is the closest to the noise curve and has the lowest AUC, followed by LE electrical. Distribution of AUCs across the cell population in retinas, shown in Fig. 6b, confirms that RCS responses were the noisiest, followed by the LE electrical responses. By replacing spikes in LE visual responses with randomly timed spikes (see Methods), we can generate a family of CCCs with various noise ratios (Fig. 6c). At a certain noise ratio, the noise-injected visual CCC matches that from electrical responses. Compared to the LE visual response with median AUC, the noise ratio was 55.3±22.5% and 78.2±6.5% for LE electrical and RCS, respectively (Fig. 6d). All RCS responses were at least 55% noisier than natural.

### Ensemble encoding

Accuracy of decoding the orientation of Landolt-C rises with increasing number of recruited cells (Fig. 7b). Since the cells were ranked by their independent decoding accuracies, the first few cells contributed to the faster rise in accuracy. In addition, beyond the first few, recruited cells started carrying redundant information, which improved accuracy with diminishing returns. Such trend is generally observed in decoding the ensemble of neural signals for applications in brain-machine interfaces (24). Notably, neither presence of photoreceptors nor stimulation type affected the decoding accuracy significantly, despite the spiking being much more stochastic under electrical stimulation, as discussed previously. Decoding accuracy also rises steeply with increasing C size until it reaches 4-5 pixels (Fig. 7b), where the accuracy flattens out because the gap in C now exceeds one pixel, and hence it is fully resolved. The asymptotic level of accuracy was determined primarily by the number of recorded cells in the retina. For example, retinas with 49 recorded cells can reach 60-80% accuracy, while only 50% can be achieved with 19 to 21 cells (Fig. 7c).

Increasing the number of stimulus presentations also increased the decoding accuracy (Fig. 7d). Therefore, to compensate for fewer cells responding in RCS retinas, more presentations are required to accumulate the same amount of information for the image decoding. To achieve 75% accuracy in decoding the orientation of letter C, LE retinas required 2-3 flashes of the image, while the RCS retinas needed 13 presentations.

## Discussion

The fact that predictive retinal models perform worse for RCS under electrical stimulation than LE retina under visual stimulation is not surprising and can be explained by increased spontaneous firing rate in the degenerate retina (25), likely due to higher uptake of retinoic acid (26). However, the fact that response of the healthy retina to electrical stimulation is much noisier than natural requires a different explanation. This might be related to the difference in mechanisms of natural and electrical activation of the photoreceptors. In natural vision, due to the rather slow phototransduction cascade, a millisecond flash causes photoreceptor hyperpolarization for tens of milliseconds (27). Under electrical stimulation, however, membrane potential is affected directly, and therefore it closely follows the electrical pulse duration (<10ms), much shorter than the natural response. Another factor might be related to the fact that our experiments were performed in the dark. Since the dark-adapted photoreceptors are depolarized, further depolarization of the terminals by electric field is quite limited, effectively restricting the dynamic range. Both factors likely contribute to the lower than natural signal-to-noise ratio when the healthy retina is stimulated electrically.

By construction, under a radially symmetric stimulus, the STA of an RGC is the first-order term in the Wiener kernel series expansion of the cell’s response function (28). Therefore, the LN model can be considered a single-filter approximation, while the CNN model can fit better due to inclusion of multiple linear filters and a better approximation of nonlinearities (29). Indeed, previous studies have shown that CNN models fit markedly better than the LN model in the salamander retina (19), which we also observed here in the healthy rat retina under visual stimulation. Surprisingly, there is little difference between the two models fitted to LE retina OFF cells stimulated electrically, indicating that the responses were predominantly single-filter (Fig. 4b). An interpretation is that OFF cell responses can be described nearly completely using only a single receptive field, which means these cells only respond to a limited subspace of stimuli. Consequentially, these RGCs may fail to respond to certain classes of spatial patterns that require subunit computation, such as null stimuli (30), where the linear filter of a cell is scaled and subtracted away from an otherwise response-inducing white noise stimulus. The disparity between LN and CNN models for RCS data suggests that even the degenerate retina retains a high degree of higher-order computation. For example, from the response to alternating grating (both ex-vivo and in-vivo), we know that nonlinear summation of subunits occurs also in the degenerate retina, but whether the number of computational subunits for each RGC matches that of the healthy retina remains unknown.

In ensemble encoding, we made two important assumptions: First, the input strength (*w* · *s*) was calculated with linear weights extracted from the binary white noise. Since pixels in the stimulus are spatiotemporally independent, the resulting trained weights are generally biased against spatiotemporally correlated stimuli, such as long straight edges and bars, drifting objects, and even natural scenes (31). Therefore, the current method leads to underestimating the accuracies in the LE retina. It is unknown whether directional sensitivity remains intact in the degenerate retina or how many nonlinear subunits exist under electrical stimulation, so the accuracy curves for the RCS retina in Fig. 7b may or may not be underestimated. Second, the letter C was placed at the same location over the 5 frames it was displayed. Normally with microsaccades, visual pattern can be displaced by ~70μm between the frames presented at 20 to 33Hz (32). Since receptive fields form a tightly packed mosaic, we assumed translational symmetry in response, i.e. no matter where the stimulus is displayed on the retina, the retinal output will carry equivalent amount of information. As a result, the current analysis assumes that the effect of the eye movements on amount of information for pattern identification is well-approximated even without moving the letter C.

Single-cell SNR played little role in ensemble encoding of the visual information. As demonstrated in Fig. 7b, all retinas had similar accuracies, even though LE RGCs under visual stimulation had far better SNR. A reason might be that better SNR as calculated might not limit the amount of information propagating downstream, if the encoded visual signals were orthogonal to the major noise eigenmodes (33). Since visual information is distributed across the retina, the more cells recruited for decoding, the higher is the accuracy. Unlike natural visual response, electrical stimulation affects the bipolar cells stronger if they reside near the electrode surface, and hence fewer RGCs were responding than in natural stimulation. To compensate for the reduced amount of visual information transmitted, the stimulus needs to be replayed multiple times. This may explain the longer time patients require to recognize letters and other patterns in clinical trials.

As in Sloan font, the gap in a Landolt-C is 1/5 of the letter size (34). For letter sizes smaller than 4 pixels, the gap is not fully resolved but encoded in some shade of grey different from the rest of the ring, which led to lower accuracy in identification. Once the gap is fully resolved, i.e. letter size greater than 5 pixels, decoding accuracy remains relatively stable. This signifies that with the pixel size used in these studies (70 μm), the limiting factor in resolution is strictly at the implant level (pixel size), but not biological (subunit size). Essentially, prosthetic vision can resolve spatial features down to the pixel size with stable accuracy, which matches our previous *in vivo* measurements (6).

In conclusion, we found that LN and CNN models matched the RGC activity elicited by subretinal electrical stimulation less accurately than that for natural responses, likely due to the weaker than natural response and higher spontaneous firing in the degenerate retina. Despite the noisier signal, visual information is still encoded across the ensemble of cells in the retina, which allows patients to perform visual discrimination tasks, albeit slower due to the reduced number of responding RGCs, compared to natural vision.

## Acknowledgements

We would like to thank Prof. Shaul Druckmann from Stanford University for advice and very helpful discussions.

Supported by the National Institutes of Health (Grants R01-EY-018608, R01-EY-027786, P30-EY-026877), the Department of Defense (Grant W81XWH-15-1-0009), Wu Tsai Institute of Neurosciences at Stanford, and Research to Prevent Blindness. Photovoltaic arrays were fabricated in the Nano@Stanford labs, which are supported by the National Science Foundation under award ECCS-1542152.

D.P. is consulting for Pixium Vision. D.P.’s patents related to retinal prosthesis are owned by Stanford University and licensed to Pixium Vision. A.S. and E.H. declare no competing financial interests.

