## Supplemental Material for "Decoding network-mediated retinal response to electrical stimulation: implications for fidelity of prosthetic vision"

*Adjusting the LN model to the data drifting*

During the MEA recording, baseline cell activity and responsivity can drift for a multitude of reasons, including fluctuations in temperature, cell health, and perfusion. In our experimental configuration for subretinal stimulation, electrical coupling between the implant and the inner retinal neurons may also change over time. Panel a in this Figure shows an example RGC activity under subretinal white noise stimulation over one hour of recording. We found that if the linear filter, i.e. the spike-triggered average (STA), is fixed after fitting to the first 10% of the recording (panel b, dotted red lines), the nonlinearity (NL) was displaced for some fragments (solid red lines) (summarized in panel c). To adjust for this drift on testing data (as opposed to training data), we included a single free parameter that can be fitted during the test time. The free parameter essentially marks the NL displacement. Without this adjustment, the model prediction and the test data can become practically uncorrelated in severe cases (panel d), while adjusted test data can restore the test accuracy close to the train accuracy.

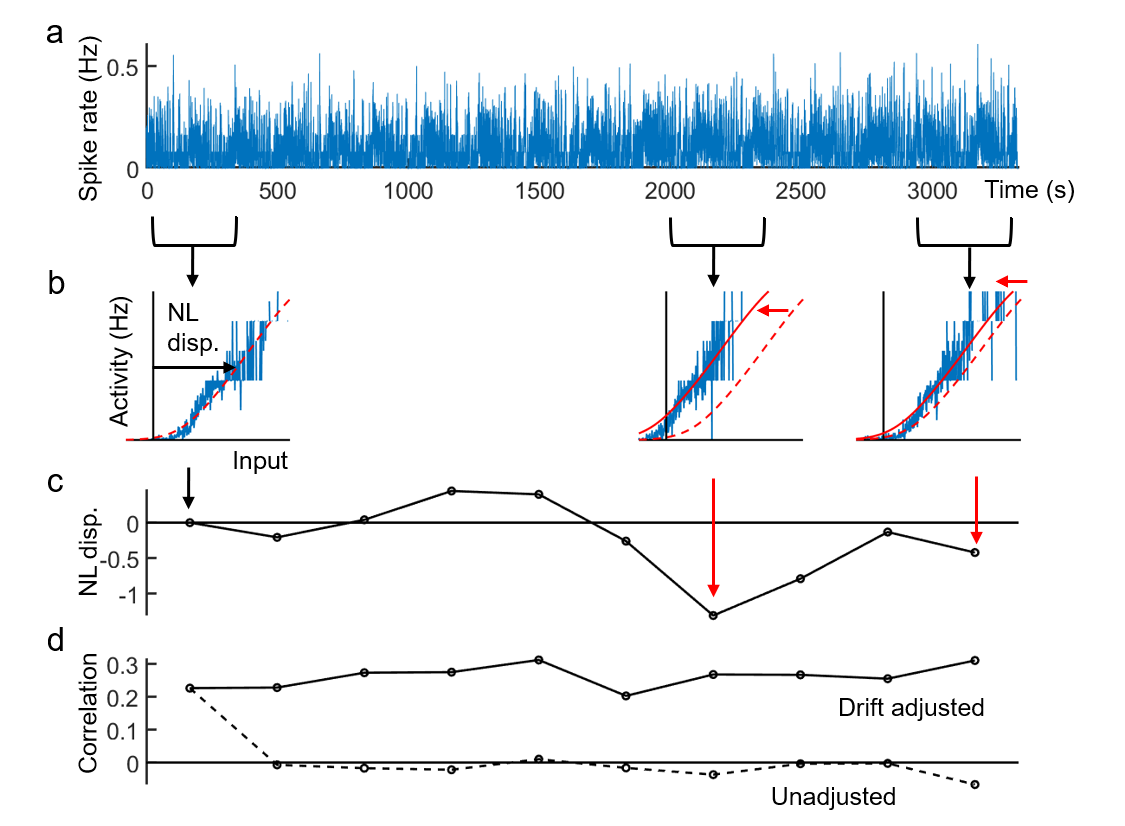
